# Parameters of Cable Theory Are Mostly Unaffected by the Geometry of Dendritic Spines

**DOI:** 10.1101/2023.08.03.551798

**Authors:** Florian Eberhardt

## Abstract

Dendritic spines are extremely small and experimentally difficult to access. Therefore, it is still uncertain whether all assumptions of basic neuroscientific theories, such as cable theory, are valid there. Previous theoretical work suggests that electroneutrality could be violated in dendritic spines. If this were true, new theories would be required. Unfortunately, these results were based on a greatly simplified model system with unrealistic ion concentrations.

Inspired by these studies, we apply Poison-Nernst-Planck (PNP) equations to study the profiles of ion concentrations and the membrane potential in dendritic spines in a physiologically relevant regime. We find that, for realistic ion concentrations and in contrast to previous results, electroneutrality is a valid assumption for all tested geometries, irrespective of size and shape. However, the surface charge causes an accumulation of counter ions and a strong electric field near the surface of the membrane in the intra- and extracellular space.

Still, a plate capacitor model accurately describes the capacitance of the membrane. Most importantly, the two cable parameters - the specific capacitance and the intracellular resistivity - are constants over a wide range of parameters. These results justify the application of models based on cable theory to dendritic spines.

## 1 Introduction

Almost all excitatory connections to pyramidal cells are made on dendritic spines [14]. Spines are therefore the first processing unit for most synaptic transmissions, and their study is essential for our understanding of synaptic function [9, 19]. Dendritic spines often have a characteristic mushroom-shaped morphology [1]. In a simplified view, they can be represented by a cylindrical spine neck and a spherical spine head [4].

The volume of a pyramidal cell spine head lies between 0.01 − 0.30 *µm*^3^. The spine neck length ranges between 0.1 − 2.21 *µm*, and the neck diameter ranges between 0.04 − 0.51 *µm* [1, 8]. The intracellular volume is bounded by the cell membrane, which consists of a lipid bilayer and other proteins. The cell membrane of neurons measures approximately 4 − 5*nm* in thickness and is negatively charged (roughly 0.02*Cm*^−2^) under physiological conditions. The surface charges are countered by a layer of positive excess ions near the cell membrane [6, 15].

Unfortunately, the limited size of dendritic spines poses a great difficulty for electrophysiological experiments, such as patch-clamp [18]. Consequently, it is still unclear whether the underlying assumptions of existing neuroscientific theories are valid there [10, 12].

Previous work suggests, that due to the high curvature of the cell membrane in spines a layer of unscreened ions extents from the membrane far into the intracellular space and violates electroneutrality in spines [3, 5]. This indicates that one of the most fundamental assumption of cable theory might not be fulfilled in small cellular compartments [10]. However, these findings have to be taken with care as the system under study was greatly simplified [2]. The results were based on simulations where only a surface charge density of 0.001 *C* · *m*^−2^ was considered, screened by 10^5^ sodium ions in the intracellular space in a sphierical volume of 4.2 *µm*^3^ [3]. This means that only a small fraction of the total charge present in spines was taken into account.

This study investigates the applicability of cable theory to dendritic spines and explores the profiles of ion concentrations and membrane potential in small neural compartments with physiological membrane charge density and ion concentrations. To achieve this, we solve the PNP-equations for geometries with radial and spherical symmetry at steady-state. Contrary to our expectations, we find that electroneutrality is a valid approximation even in very small compartments. While a plate capacitor model cannot account for the exact profiles of membrane potential and ion concentrations, it accurately describes the membrane capacitance. Cable parameters, including capacitance and intracellular resistivity, are largely unaffected by spine morphology and remain constant over a wide range of parameters. We introduce a correction factor based on spherical or cylindrical geometry only for very small geometries with radii smaller than 20 *nm*. These findings justify the application of cable theory-based models to dendritic spines.

## 2 Methods

### PNP-equations

The Nernst-Planck equation in 3D cartesian coordinates is written as

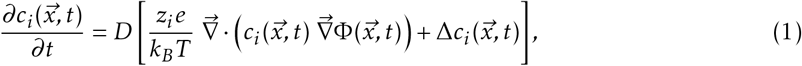

and Poisson’s equation is written as

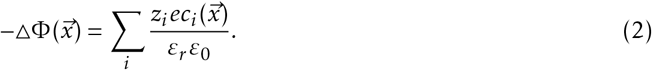

To simplify the notation further, we define the charge density *ρ* as

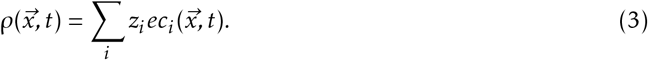

Above, *c*_*i*_ denotes the particle density of ion-species *i*, Φ denotes the electric potential, *D* denotes the diffusion constant, *z*_*i*_ denotes the charge number, *e* denotes the elementary charge, *k*_*B*_ denotes the Boltzmann constant, *T* denotes the temperature, *ε*_*r*_ *ε*_0_ denotes the permittivity, and 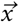 and *t* denote the variables for space and time.

### Steady-state equations for radial and spherical symmetry

We study systems with spherical and radial symmetry in steady-state, which implies that 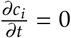. In the case of spherical symmetry, equation 1 can be simplified to

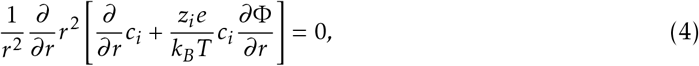

where *r* denotes the radial distance to the center of the sphere. The term in square brackets describes the radial current *j*_*r*_ . From equation 4, we can conclude that *r*^2^*j*_*r*_ = *const*.”

Next, we assume that the sum of all currents across the membrane is zero for each individual ion species, meaning that there is no net flow of ions through the membrane at *r* = *R*. This implies that *j*_*r*_ (*R*) = 0. However, as *r*^2^*j*_*r*_ is constant, it follows that the radial current *j*_*r*_ is zero everywhere. Therefore, we can further simplify equation 4 to

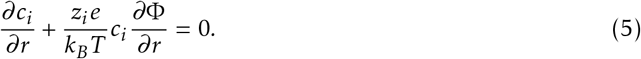

The exact same equation can be derived for cylindrical symmetry when the additional assumption is made that any current flowing along the main axis of the cylinder, *j*_*z*_, is independent of *z*, i.e., 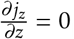.

Next, we simplify Poisson’s equation (equation 2). In spherical symmetry, Poisson’s equation can be written as

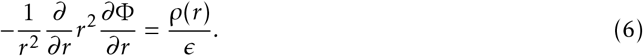

From here, integration leads to

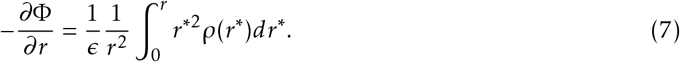

In cylindrical symmetry (here *r* denotes the Euclidean distance to the central axis of the cylinder), Poisson’s equation can be written as

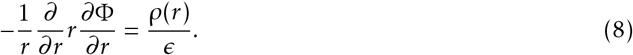

By assuming that there is no dependence on the *z*-coordinate, i.e., 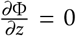, the equation can be integrated, which leads to

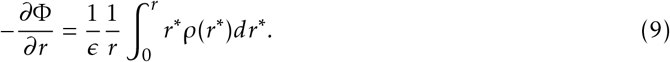

### Boundary Conditions

In both cases, spherical and polar symmetry, the electric potential and the electric field at a radius *r* = *r*^∗^ only depend on charges inside a sphere or a cylinder of this radius *r*^∗^. Charges more distant from the central point or central axis do not contribute. Consequently, if the charge density at the center of the sphere or disk (*P*_*A*_, Fig. 1) were to sum up to zero exactly (*ρ*(0) = 0), no potential would build up. Therefore, we assume only that at some distant point *P*_*F*_ (Fig. 1), the bulk ion concentrations sum up to exactly zero. Now, the central point *P*_*A*_ is in electro-chemical equilibrium with this distant point *P*_*F*_. Using the following equations, the ion densities and the electric potential can be weakly perturbed from the zero-charge (*ρ*(0) = 0) state at *P*_*A*_ by assuming that Φ(*r* = 0) = Φ^*F*^ + *δ*Φ. Using the equilibrium assumption between *P*_*A*_ and *P*_*F*_, this leads to

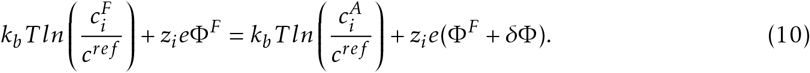

By solving this equation for *c*_*i*_ one arrives at

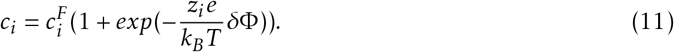

We make the same assumptions for the extracellular points *P*_*E*_ and *P*_*D*_, respectively. Having introduced these parameters, we can state the boundary conditions for the system. Below, we consider all quantities to be only functions of the radial distance *r*. The boundary conditions are:

**Figure 1:**
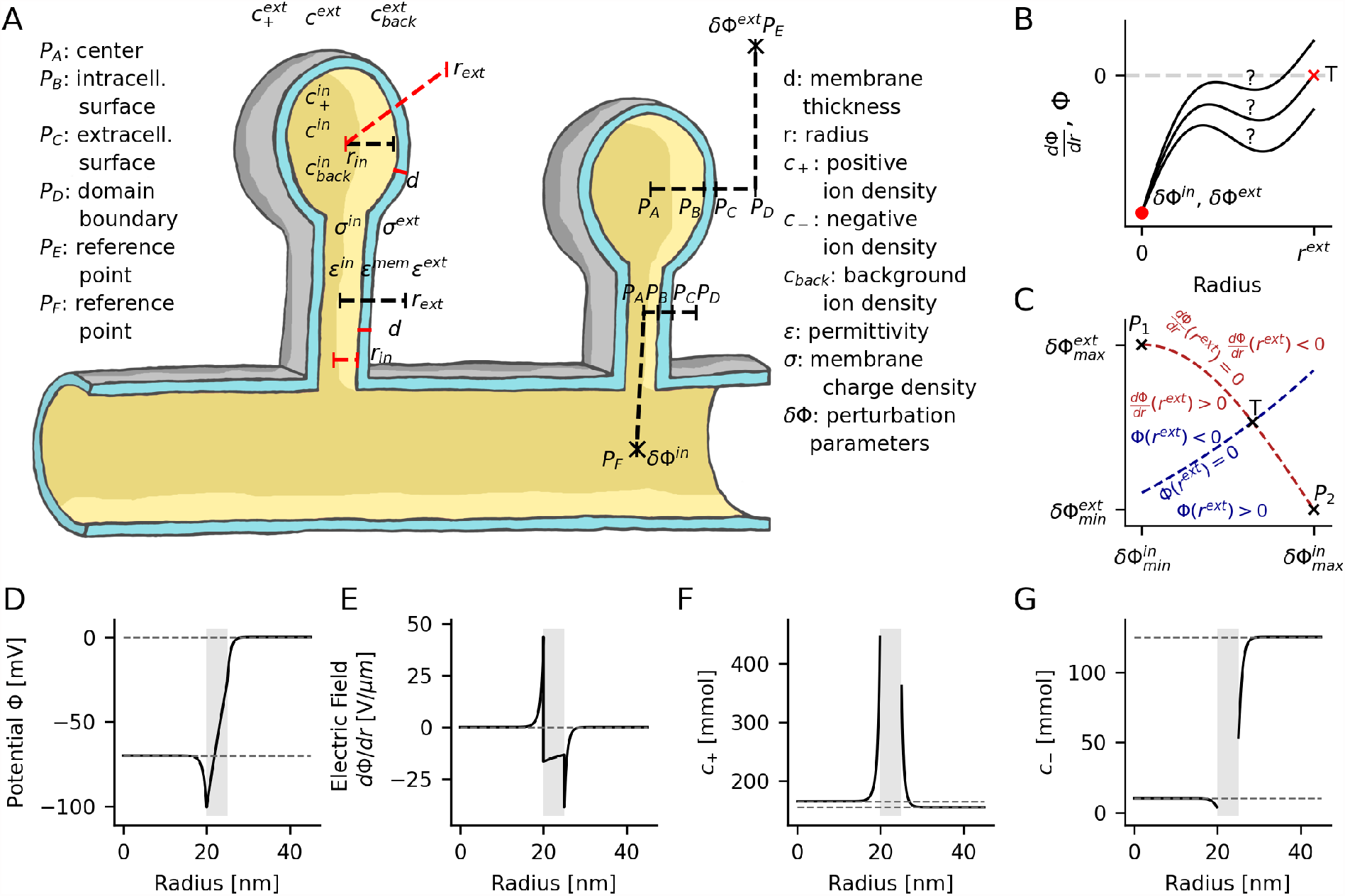
Solution of the PNP-equations for radial and spherical symmetry at steady-state. A) Notation for variables and parameters used in this study. B and C) A shooting method is used to solve the PNP-equations as an initial value problem. The electrode potential at the center of the domains (*P*_*A*_) and at the extracellular surface of the membrane (*P*_*C*_) are perturbed with the parameters *δ*Φ^*in/ext*^ until a solution that fulfills all boundary conditions is found. D-G) An example solution of a domain with polar (cylindrical) symmetry for an intracellular radius of 20 *nm* is shown. The center of the domain corresponds to the radius *r* = 0 *nm*. The gray region indicates the cell membrane, with the intracellular space on the left side and the extracellular space on the right side of the membrane. Fixed negative background charges, which are 155 *mmol* in the intracellular space and 30 *mmol* in the extracellular space, are not shown.

#### A) central point

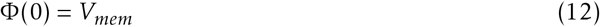

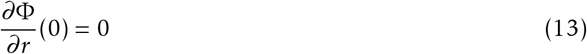

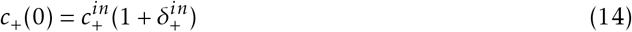

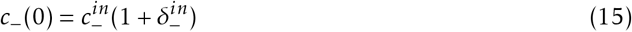

where

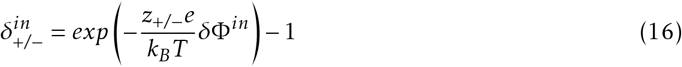

*V*_*mem*_ denotes the membrane potential, which is assumed to be − 70 *mV* in the resting state. Due to the spherical or polar symmetry, there is no electric field in the radial direction at the central point. Ion concentrations at the center are the physiological intracellular ion concentrations, weakly perturbed by a small perturbation voltage *δ*Φ^*in*^.

#### B) Intracellular membrane surface

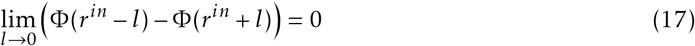

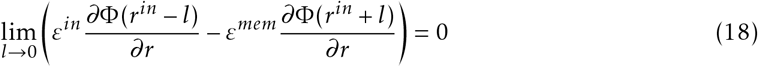

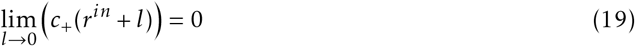

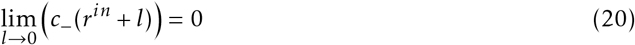

The electric potential is continuous, and the jump in the electric field is given by the change in permittivity between the intracellular space and the membrane. Additionally, the number of free ions inside the membrane is zero (or negligible).

#### c) Extracellular membrane surface

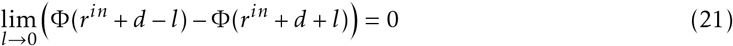

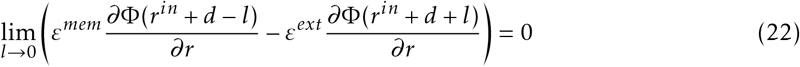

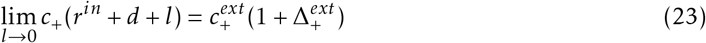

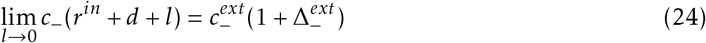

where

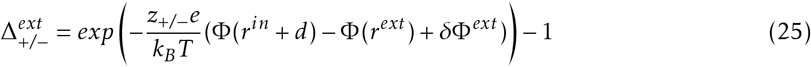

Again, the electric potential is continuous. The jump in the electric field is given by the change in the permittivity between extracellular space and membrane. Point *P*_*D*_ (Fig. 1) is in electrochemical equilibrium with some distant point *P*_*E*_.”

#### D) Exterior boundary

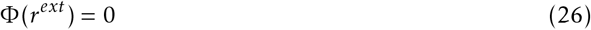

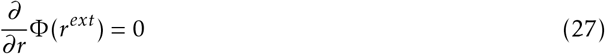

The voltage difference between central point *P*_*A*_ and exterior boundary *P*_*D*_ is determined by the membrane potential. The system is globally electroneutral; hence, there is no net charge at the exterior boundary, and the electric field is zero there.

### Numercial implementation

The perturbation parameters and the previous boundary conditions allow us to formulate the problem as an initial value problem. Therefor, the perturbation voltages *δ*Φ^*ext*^ and *δ*Φ^*in*^ are fixed, and the system is solved starting from the center using an explicit Runge-Kutta algorithm. Then, the perturbation voltages are optimized using a shooting method (Fig. B, C) until the boundary conditions at point *P*_*D*_ are fulfilled.

In the following, we will derive the Runge-Kutta solver for our problem. By inserting Eq. (7) into Eq. (5), we find in the case of a single ion species

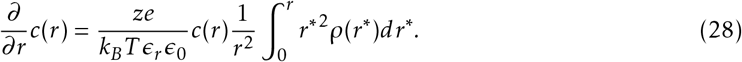

In the case of multiple ion-species the previous equation can be rewritten as

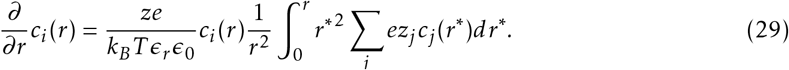

To solve eq. 29 with an explicit Runge-Kutta algorithm, we first need to estimate the term under the integral in this equation. Under the assumption, that the concentrations *c*_*i*_ (*r*) are known for 0 < *r* < *r*_*n*_, one can compute *c*_*i*_ (*r*_*n*+1_) by

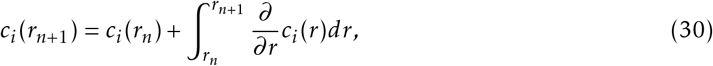

where *r*_*n*+1_ = *r*_*n*_ + *h*. Together with (29) one finds

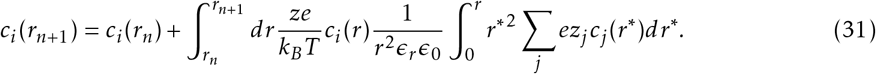

The second integral can be split up into two terms

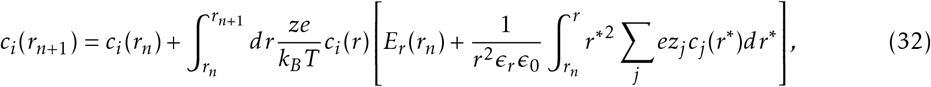

where *E*_*r*_ (*r*_*n*_) denotes the electric field at radius *r* = *r*_*n*_. After approximating the term under the inner integral with the constant term of the Taylor-expansion one arrives at

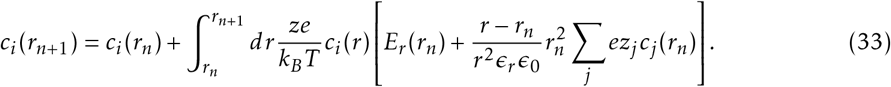

Finally we define

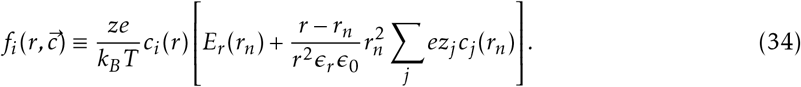

After setting the initial values at *r* = 0, the concentrations can be computed using the following scheme of an explicit RK4 algorithm, as presented in [13]:”

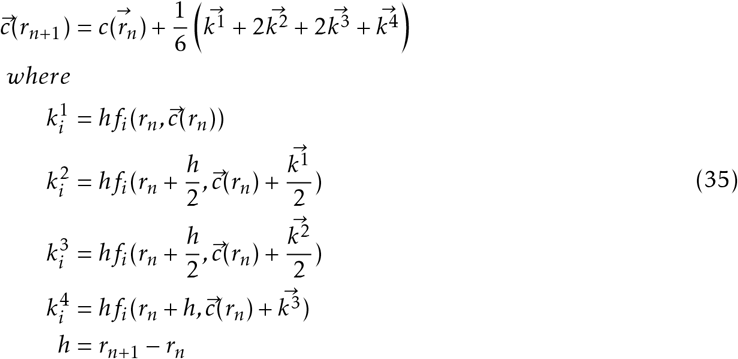

The function *f*_*i*_ can also be derived for polar symmetry, this leads to

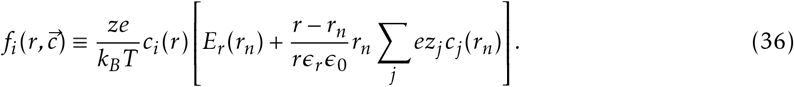

To ensure the accuracy of the numerical solution, we use multi-precision floating-point numbers with up to 1000 digits.

### 3 Results

In the following, we refer to the value set for the Dirichlet boundary condition of the variable Φ at the center of the domain as the electrode potential *V*_*m*_. This value can be compared to the membrane potential, which can be measured between two electrodes located in the intra- and extracellular space. The term electric potential denotes the solution of the variable Φ(*r*) and is a function of the independent variable *r*, representing the radial distance to the center.

### Solution of PNP-equation

We solve the PNP equations in domains with spherical and radial symmetry at steady-state. Each domain consists of an intracellular space with a radius of *R*_*in*_, a membrane with a thickness of *d* = 5*nm*, and an extracellular space. The external boundary of the simulated domain is located at a radius of *R*_*ext*_ = 2*R*_*in*_+*d* (Fig. 1A). The radius *R*_*in*_ varies between 10*nm* and 80*nm* in domains with radial symmetry (cylindrical spine necks) and between 10*nm* and 160*nm* for spherical domains (spine heads). This includes the lower boundary of spine sizes [8]. For each geometry, we solve the PNP-equations for four different electrode potentials *V*_*m*_ ranging from − 70*mV* to +35 *mV* . Thereby, we compute the electric potential Φ, the electric field *d*Φ*/dr*, and the concentration of positive *c*_+_ and negative *c*_−_ ions as a function of the radial distance *r* to the center of the cylindrical and spherical domains (Fig. 1D-G).

The profiles of ion concentrations and electric potential are qualitatively similar in all tested geometries and for all electrode potentials. Only the voltage drop and electric field inside the cell membrane depend on the chosen electrode potential. An example of a cylindrical geometry with *R*_*in*_ = 20 *nm* and *V*_*m*_ = −70 *mV* is shown in Figure 1D-G. Starting from the center of the intracellular space at *r* = 0 *nm*, the electric potential Φ is initially constant and then decreases near the membrane. Inside the membrane, the electric potential linearly increases by approximately 70 *mV* . Outside the membrane in the extracellular space, Φ rapidly converges to 0 *mV* and remains constant until the external boundary at *r* = *R*_*ext*_ is reached (Fig. 1D).

The electric field is approximately constant inside the membrane, with its strength and direction depending on the electrode potential *V*_*m*_. In both the intra- and extracellular space, the electric field increases as a function of *r*, indicating an excess of positive charges on both sides of the membrane. The existence of positive excess charges is also confirmed by the ion concentration profiles. Regardless of the electrode potential *V*_*m*_, there is always a strong accumulation of positive charges and a depletion of negative charges at the membrane.

### The membrane charge maintains the shape of the concentration profiles

To better understand the found solutions, we compared the distribution of excess charges *c*_*excess*_ = *c*_+_− *c*_−_ for different electrode potentials *V*_*m*_ and various radii (an example of a spherical geometry with *R*_*in*_ = 80 *nm* is shown in Fig. 2A). Regardless of the geometry and the applied electrode potential, there is always a high concentration of positive excess charges at the membrane surface. Moreover, the peak concentration of the excess charge depends only weakly on the electrode potential. For *V*_*m*_ = +35 *mV*, there are slightly more positive excess charges on the intracellular side than for *V*_*m*_ = − 70 *mV* . The difference is weak compared to the absolute concentration of the excess charges in all studied cases.

**Figure 2:**
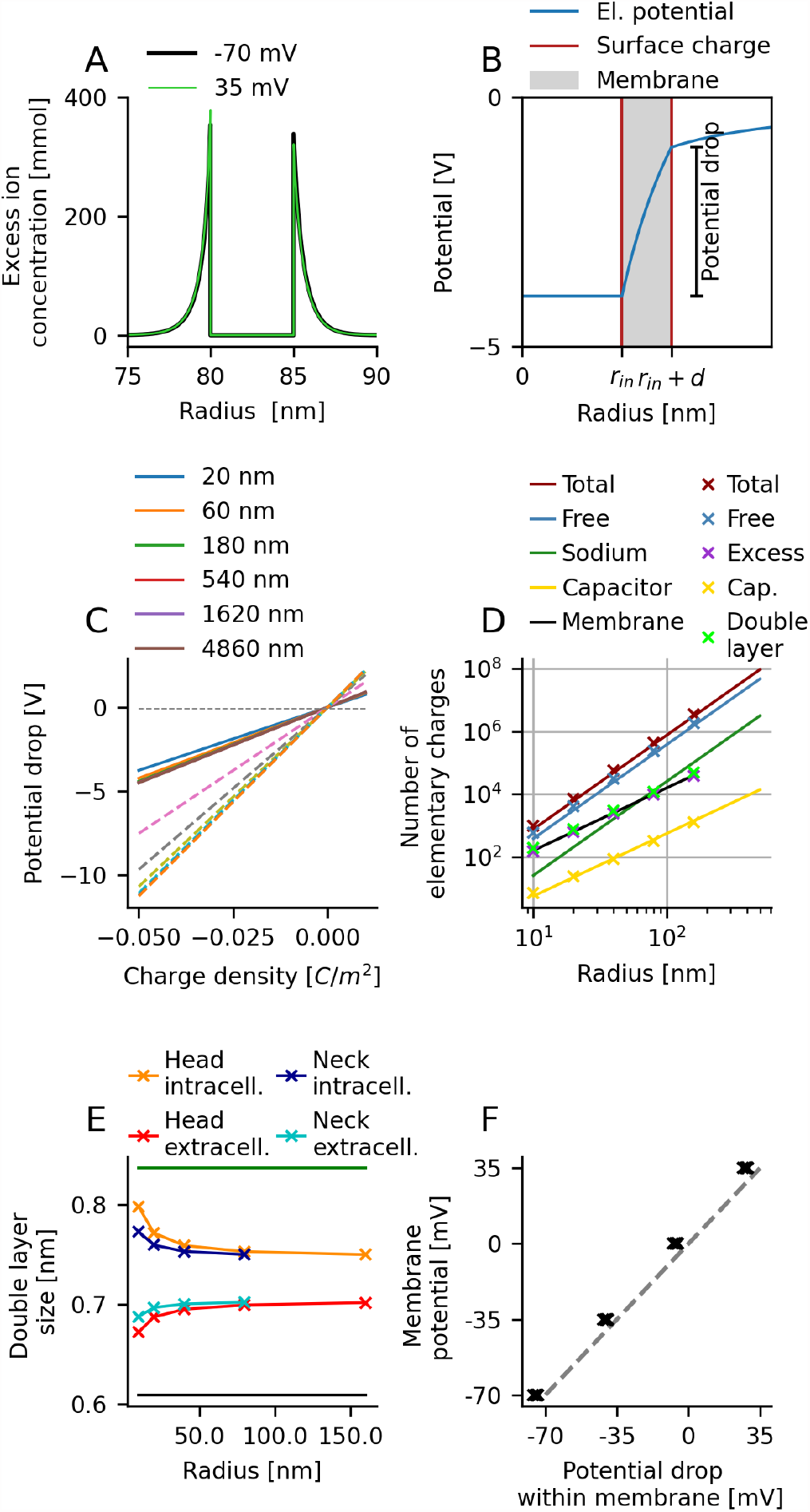
Electroneutrality is a valid assumption in dendritic spines. A) The concentrations of excess ions correspond to the density of unscreened ions (*c*_+_ *c*_−_). Excess charges only occur close to the membrane surface. On both sides of the membrane, the excess charge is strongly positive, irrespective of the electrode potential. B) Outline of a purely hypothetical scenario where the intracellular and extracellular space, and the interior of the membrane, are void of charges. The membrane carries a negative surface charge. This leads to a potential drop across the membrane. C) In the same system as in (B), a membrane charge of − 0.02 *C/m*^2^ evokes a potential drop of several volts across the membrane. This is much stronger than the physiological resting potential of − 70 *mV* . D) Excess charges cannot be explained by the charge on the membrane capacitor. The number of excess charges is comparable to the number of membrane charges but an order of magnitude lower than the total number of free charges in spines. E) The distance at which the excess concentration drops to 1*/e* compared to its peak value is at the size of the Debye length. The system can, therefore, be assumed as electro-neutral. The black line includes background charges to estimate the Debye length. The green line ignores the background charges when estimating the Debye length. F) The excess charges occur only close to the membrane. The potential drop in the excess charge layer in the intracellular and the extracellular space almost compensates. Therefore, the voltage drop inside the membrane is comparable to the electrode potential *V*_*m*_.

This suggests that the negative surface charge of the membrane strongly accumulates positive charges at the membrane surface. To further investigate the effect of the membrane charges, we studied a hypothetical system that only carries negative membrane charges and is free of charges in the intracellular and extracellular space. Consequently, the membrane charges remain unscreened, resulting in a voltage drop of several volts across the membrane (Fig. 2B). The found values vary across different geometries from roughly 2 *V* for a membrane thickness of 4*nm* (Fig. 2C, solid line) up to 7*V* for a membrane thickness of 10 *nm* (dashed line) at a charge density of −0.02 *C/m*^2^. This found hypothetical voltage drop across the membrane is always significantly higher than the physiological membrane potential.

Next, we compared the total number of excess charges in the intracellular space (purple marker in Fig. 2D) to the number of surface charges on the intracellular side of the membrane (black line in Fig. 2D) and found that these numbers agree well. Therefore, it can be concluded that most excess charges compensate the surface charge and do not contribute to the membrane potential.

Nevertheless, a difference in the membrane potential still results in a small difference in excess charges (Fig. 2A). Therefore, we computed the difference in the total number of excess charges in the intracellular space between a membrane potential of *V*_*m*_ = 0 *mV* and *V*_*m*_ = −70 *mV* . We compared this number to the theoretically predicted number of charges needed to charge a plate capacitor with size *A* = 4*πr*^2^ to *V*_*cap*_ = −70 *mV* . Again, we find that the numbers match well. Thus, the difference between the excess charge concentrations (green and black line in Fig. 2A) can be explained by a discharging and charging of the membrane capacitance. In summary, we conclude that ion-concentration profiles are mostly maintained by the negative surface charge of the membrane and are only weakly affected by the membrane potential.

### Electro-neutrality is a valid assumption in dendritic spines

Previously, it was proposed that the high curvature of the membrane in dendritic spines perturbs the concentration profiles of excess charges and breaks electroneutrality in spines [10, 3, 5]. However, these results were based on a model system in which only a reduced number of negative charges was fixed on the surface of a sphere. The identical number of positive charges was then placed to freely move in the intracellular space. The electric potential and ion concentrations were computed by solving PNP equations. However, the total number of charges present in this system is comparable to the number of charges needed to charge a membrane capacitor (yellow line in Fig. 2D). Hence, this model considers only a small fraction of all ions present in spines. The total number of charges is much higher in real spines (blue and red lines in Fig. 2D).

To test whether electro-neutrality is also broken under more realistic conditions, we examine the concentration of excess charges at physiological ion concentrations. In theory, an electrolyte is electro-neutral if there are no electrostatic forces acting beyond the Debye length, which is below 1 *nm* in neural tissue. Within one Debye length, the electric potential decreases in magnitude by a factor of 1*/e*. Therefore, we measure the distance from the membrane where the electric potential drops by 1*/e*. We find that this distance ranges between 0.67 *nm* and 0.80 *nm* for all studied domains. The theoretically estimated Debye length, including fixed background charges (green line in Fig. 2E), and excluding background charges (black line in Fig. 2E) is of the same size. It can be concluded that electro-neutrality is an excellent approximation even in very small neural compartments.

There is still a small but visible difference between the size of the membrane layer on the intracellular and extracellular side (Fig. 1E). The difference can be explained by the sign of the curvature. On the convex intracellular surface, the layer of excess charges extends further into the intracellular space compared to the extracellular side where the surface is concave. This difference leads to a larger potential drop in the intracellular space compared to the extracellular space. In turn, this can lower the value of the potential drop inside the membrane below the electrode potential.

To test if there is a relevant a geometrical effect on the electric potential, we compared the potential drop inside the cell membrane with the applied electrode potential. We found that the potential drop within the membrane is only slightly smaller than the electrode potential. This indicates that the potential drop in the intracellular and the extracellular space almost compensate. Moreover, there is only a negligible effect by the geometry, as the black crosses overlay in Fig. 2F.

In summary, the geometry and the electrode potential only weakly affect the distribution of ions and the electric potential in dendritic spines.

### Effect on cable parameters are irrelevant for most spines

The ion concentrations and electric potential are only weakly affected by the electrode potential and spine geometry, and electro-neutrality is a valid assumption in spines. In the following section, we will investigate whether other assumptions of cable theory may still be violated.

The membrane potential in the spine head can differ by several millivolts compared to its parent dendrite [7]. In such cases, an electric current flows through the spine neck. In cable theory, current flow is assumed to be exclusively axial. To verify this assumption for thin spine necks, we solved the PNP-equations for different electrode potentials of *V*_*m*_ = − 70*mV*, 35*mV*, 0*mV*, and 35*mV* . We then computed the difference between the electric potentials of two solutions, e.g., *V*_*dif f*_ (*r*) = Φ(*r*)+35 Φ(*r*) 70. In Fig. 3A, the dotted lines indicate the solutions of the PNP-equations, while the colored lines indicate the difference between the electric potentials at *V*_*m*_ = − 70*mV* and depolarized states *V*_*m*_ > − 70*mV* .

**Figure 3:**
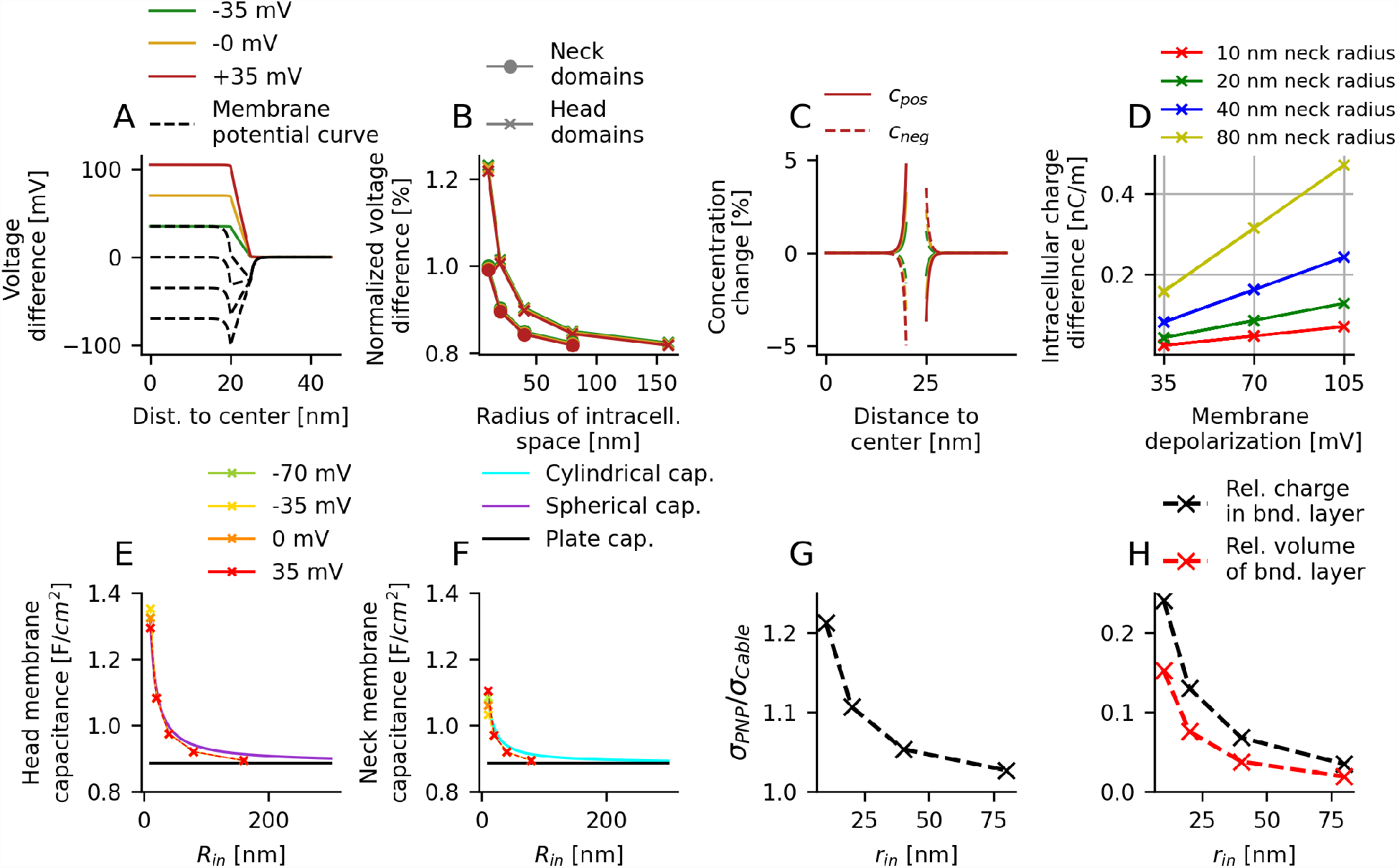
Parameters of cable theory in dendritic spines. A) The dashed lines indicate the electric potentials as found by the PNP-equations. The solid lines indicate the difference between the electric potentials at *V*_*m*_ *>* − 70*mV* with the electric potential at *V*_*m*_ = − 70*mV* . In radial direction, the difference between the two electric potentials is almost constant. B) The maximum variation along the solid lines in A) in the intracellular space is always significantly smaller than the difference between the two electrode potentials. This indicates that radial current flow is only weak when the membrane potential is changing. C) The difference between the two concentration profiles corresponding to the same solutions used to compute the solid lines in A) is shown. The ion concentrations are only affected at the membrane and reflect charging and discharging of the membrane capacitance. D) In depolarized states (*V*_*m*_ *>* − 70*mV*) the total charge in the intracellular space increases linearly with the depolarization. The membrane behaves as an ideal capacitance. E) The capacitance increases in domains with spherical symmetry for small radii. The relative change can be explained by the spherical geometry. F) Changes of the capacitance in domains with polar symmetry can be explained by a cylindrical capacitor. In larger domains, the capacitance is well approximated by a plate capacitor in E) and F). G) The normalized conductivity along a cylinder increases for small radii. This is independent of the membrane voltage and can be explained by the increasing relative volume of the boundary layer, as seen in H). In larger domains, this effect vanishes.

We find that *V*_*dif f*_ is almost constant in the intracellular (or extracellular) space (Fig. 3A). When we compare the maximum variation of *V*_*Dif f*_ within the intracellular space with the difference between the two respective membrane potentials, we find that the voltage difference in radial direction hardly exceeds 1 % of the axial difference (Fig. 3B). We conclude that in steady state, current flow in a thin cylinder is well-assumed as axial current.

Next, we want to know whether the membrane in spines behaves as an ideal capacitor. In theory, the total capacitance *C*_*m*_ of a neuron’s membrane is directly proportional to its surface *A*, and can be computed as *C*_*m*_ = *c*_*m*_*A*. The specific membrane capacitance *c*_*m*_ is roughly 1 *µF/cm*^2^. The capacitor voltage depends linearly on the charge *Q*, which is located on the two surfaces. To charge the membrane capacitance to a voltage *V*_*cap*_, a charge *Q* = *CV*_*cap*_ is needed. When we compare the concentration profiles of *c*_+_ and *c*_*−*_ for the electrode potentials *V*_*m*_ = − 35, 0, +35*mV* with the ion concentrations for *V*_*m*_ = − 70*mV*, we find that only ions in close proximity to the membrane are relocated. The difference between the concentrations is up to 5 % in the membrane layers and 0 elsewhere (Fig. 3C). The membrane surface can therefore be considered as the capacitor surface. To further test the linear dependence between voltage and charge, we compute the total charge in the intracellular space. Then we compute the difference between the total charge at different electrode potentials. We find that the difference is a linear function of the membrane potential, independent of the size of the domain, and as expected from an ideal capacitor (Fig. 3D).

To further investigate whether the total capacitance *C*_*m*_ is a linear function of the surface *A*, as expected for a plate capacitor, we test if the specific capacitance *c*_*m*_ = *C*_*m*_*/A* is independent of the radius of the geometry. We compute the specific membrane capacitance as a function of the intracellular radius 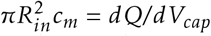. We find that for very small radii *R*_*in*_ = 10 nm, 20 nm in cylindrical geometries (Fig. 1F) and *R*_*in*_ = 10 nm in spherical geometries (Fig. 3E), the relative change is higher than 10 %. In both cases, the deviations from a plate capacitor can be explained by geometrical corrections as spherical or cylindrical capacitors. For larger geometries comparable to most dendritic spines, the capacitance is well approximated by a plate capacitor.

Finally, we ask if the electric resistance of a spine neck is affected by the neck’s diameter. In cable theory, the cytosol is modeled as an ohmic resistor. The total electric resistance *R*_*e*_ of a cylinder with cross-section *A* and length *l* is *R*_*e*_ = *r*_*e*_*l/A*, where *r*_*e*_ is the intracellular resistivity. The resistivity linearly depends on the ion concentration and is the inverse of the conductivity *σ*. However, inside the boundary layer (size measured in Fig. 2E), the ion concentrations are elevated (Fig. 1F, G, and Fig. 2A). In small neural compartments with a high surface-to-volume ratio, like dendritic spines, the impact of the boundary layer could become dominant. Indeed, for very small radial geometries, the conductivity increases (Fig. 3G). This can be well explained by the relative size and the relative ion content of the boundary layer (Fig. 3H). However, this effect exceeds 10% only for radii less than or equal to 20 nm.

In summary, cable parameters of geometries comparable to dendritic spines, that are usually larger than 20 nm are only weakly affected.

## 4 Discussion

Electro-neutrality is a basic assumption of traditional neuroscientific theories, such as cable theory, to model the electrical function of neurons. However, it has been proposed that electroneutrality could be violated in small neuronal compartments with high membrane curvature, comparable to those found in dendritic spines [3, 5, 10]. The studies mentioned, however, significantly underestimated the ion concentrations and ignored the extracellular space and the membrane with its surface charges. In particular, [3] studied a dielectric ball with a radius of 1*µm*, a negative surface charge of 0.001*Cm*^−2^ density, and 0.4 *mmol* concentration of positive ions inside the volume. However, under physiological conditions, the surface charge is roughly 0.02*Cm*^−2^ and the charge density is up to 300 *mmol* of charges in the intracellular space.

To test whether electro-neutrality is also broken under more realistic conditions, we investigate the ion concentrations and electric potential in spherical spine heads and cylindrical spine necks. We study a system with physiological ion concentrations and surface charge density, and we include the extracellular space in our model systems. For this purpose, we present a numerical method to solve the PNP equations in radial and spherical symmetry at steady state for an arbitrary composition of the electrolyte and realistic ion concentrations. Previous studies have investigated the profile of the membrane potential for flat membranes [16], but not in spherical or polar symmetry.

We find that the membrane charge causes an accumulation of excess ions on both sides of the membrane. However, contrary to what was predicted, the size of this layer of excess ions depends very weakly on the curvature of the membrane. On the intracellular and extracellular side, the size of the double layer is comparable to the Debye length. Therefore, it can be assumed that electro-neutrality is a valid assumption in dendritic spines, unlike what was proposed previously. We want to point out that the charge density in real neurons is often asymmetric [6, 17]. However, this would not affect the Debye length or the main results of this study.

Our findings contribute to the ongoing discussion on how to mathematically model dendritic spines. We quantitatively confirm the arguments of Barbour [2] on electroneutrality in small neural compartments. However, as pointed out by Holcman and Yuste [11], even in the case of electroneutrality, PNP-equations could be more suitable to model ionic currents in spines than cable theory. Moreover, classical definitions of parameters like capacitance might not be applicable to spines. To further explore this idea, we tested the validity of various assumptions of cable theory in small neural compartments.

Most importantly, we found that the membrane can be considered an ideal capacitor (Fig. 3C, D), and the conductivity is proportional to the cross-section (Fig. 3H). Only in geometries smaller than most spines did we find deviations in capacitance and resistivity. Based on these results, spines have the smallest possible size to avoid measurable geometrical effects. For physiologically relevant regimes, it is a reasonable assumption to treat cable parameters such as capacitance and resistance as constants.

In summary, the assumptions related to cable theory that were tested in this study are also valid in small neuronal compartments such as dendritic spines. However, we did not test whether the ion concentrations themselves remain constant. Previous studies have found that ion concentrations are likely to change in spines during synaptic input [12]. In combination with the results presented here, we propose using models based on cable theory with variable ion concentrations to accurately model dendritic spines.

## Acknowledgements

I am deeply grateful to my professor and supervisor, Prof. Dr. Andreas V. M. Herz, for his guidance and support throughout my doctoral studies. Furthermore, I would like to acknowledge the generous support of the Ludwig-Maximilians Universität, the German Federal Ministry of Education and Research, and the Bernstein Center for Computational Neuroscience Munich, who provided the funding necessary to carry out this research.

